# The roles of standing genetic variation and evolutionary history in determining the evolvability of anti-predator strategies

**DOI:** 10.1101/002493

**Authors:** Daniel R O’Donnell, Jordan A Fish, Abhijna Parigi, Ian Dworkin, Aaron P Wagner

**Affiliations:** Center for Microbial Ecology, Michigan State University, East Lansing, MI 48824, USA; Program in Ecology, Evolutionary Biology, and Behavior, Michigan State University, East Lansing, MI 48824, USA; Department of Zoology, Michigan State University, East Lansing, MI 48824, USA; BEACON Center for the Study of Evolution in Action, Michigan State University, East Lansing, MI 48824, USA

## Abstract

Standing genetic variation and the historical environment in which that variation arises (evolutionary history) are both potentially significant determinants of a population’s capacity for evolutionary response to a changing environment. We evaluated the relative importance of these two factors in influencing the evolutionary trajectories in the face of sudden environmental change. We used the open-ended digital evolution software Avida to examine how historic exposure to predation pressures, different levels of genetic variation, and combinations of the two, impact anti-predator strategies and competitive abilities evolved in the face of threats from new, invasive, predator populations. We show that while standing genetic variation plays some role in determining evolutionary responses, evolutionary history has the greater influence on a population’s capacity to evolve effective anti-predator traits. This adaptability likely reflects the relative ease of repurposing existing, relevant genes and traits, and the broader potential value of the generation and maintenance of adaptively flexible traits in evolving populations.

## Introduction

The diversity and complexity of any biological system reflects past evolutionary responses to environmental conditions, as much as contemporary ones. Underlying those responses are a number of extrinsic and intrinsic factors influencing populations’ capacity to evolve responses through the generation or utilization of genetic variation [1,2]. Because evolutionary responses build on available heritable variation, mechanisms that influence the generation and maintenance of variation [3] can strongly shape the evolutionary trajectories of populations [4]. Among the potential factors involved, standing genetic variation (SGV) and evolutionary history (EH) are likely to be significant determinants of adaptability to novel environments [5]. Accordingly, understanding how these critical factors either promote or constrain population evolutionary potential provides insight into the realized pathways that led to historic evolutionary outcomes, as well as those that will shape future populations.

Standing genetic variation is the presence of alternate forms of a gene (alleles) at a given locus [5] in a population. While an allele may be mildly deleterious or confer no fitness advantage over other forms under a given set of environmental conditions [6], that allele may become beneficial if the environment changes. As selection can act only on available variation, SGV provides a potential means for more rapid adaptive evolution [7] compared with the *de novo* mutations [5,8], particularly if environmental conditions change.

In addition to SGV, a population’s historical selection environment (i.e. evolutionary history) may play a strong role in determining the speed and the extent to which populations can adapt to directional environmental change [9,10]. In particular, EH will have influenced the genetic variation and genetic architecture of traits in contemporary populations. If changes in the environment alter the strength of selection on a trait, populations with an evolutionary history of adaptation to similar pressures may be mutationally “closer” to the discovery of new [11], or rediscovery of historic beneficial traits [12].

With the contemporary rise of experimental evolution as a means of testing evolutionary and ecological hypotheses [13,14], a valuable and untapped opportunity now exists to elucidate the roles SGV and EH have played in determining historic, realized rates of adaptive evolution. Furthermore, while SGV is known to be an important determinant of the speed of evolution [5], it is less clear how levels of SGV and EH, alone or in combination, impact the overall evolutionary potential of populations. Such an understanding could allow population geneticists to evaluate how SGV and EH may contribute to or constrain the future evolvability of populations, particularly in human-modified environments [1, 15, 16]

To evaluate the individual and interactive effects of SGV and EH on evolutionary outcomes, we measured their relative and independent contributions toward evolutionary potential in populations using the digital evolution software Avida [17]. Digital evolution experiments carry several significant advantages for addressing evolutionary questions requiring systematic manipulation and highly controlled environments. Among these, generation times are rapid, the experimenter has full control over initial environmental conditions, and detailed genetic, demographic, and behavioral trait data can be recorded perfectly. Organisms in Avida can also engage in ecological competition and other complex interactions, and the system can allow for co-evolution with predators in predator-prey systems [18,19,20]. Furthermore, unlike in evolutionary models and evolutionary simulations that impose artificial selection via explicit selection functions, Avida uniquely allows for unrestricted, unsupervised, and non-deterministic evolution via natural selection [17].

Predation is an ecologically important agent of selection [21,22] as demonstrated by the diverse array of prey defenses that have evolved [23,24,25,26]. Accordingly, we used Avida to test whether historic exposure to predation influenced how prey populations responded to pressures from new invasive predators. We then further examined which factors (SGV, EH, and their interaction) were important in determining the future defensive and competitive abilities of those populations. While both factors played significant roles, we show that evolutionary history has the stronger influence on evolutionary potential, with the strength of SGV’s role being contingent on a population’s evolutionary history.

## Materials and Methods

### Avida

The Avida digital evolution platform is a tool for conducting evolutionary experiments on populations of self-replicating computer programs, termed digital organisms [17]. Digital organisms are composed of a set of instructions constituting their “genome”. Organisms execute genome instructions in order to perform actions such as processing information, and interacting with their environment, and for reproduction. Additionally, predefined combinations of instructions allow organisms to consume resources from the simulated environment. Sufficient consumption of resources, to a level defined by the experimenter, is a prerequisite for organisms to copy their genomes and divide (i.e. reproduce). During the copy process, there is an experimenter-defined probability that genomic instructions will be replaced with a different instruction (substitution mutation) randomly selected from the full set of all 60 possible instructions (instruction set files are in the skel directory in the GitHub repository [https://github.com/fishjord/avida_predation_scripts]).

Separately, there are also set probabilities for the copy of an instruction to fail (creating a deletion mutation) or for a chance insertion of a new instruction into the copy’s genome. In real-time, generation times in Avida are typically only a few seconds, and experimental environments are fully definable.

#### Environment

We ran all Avida trials in a 251 × 251 bounded grid-cell environment. Each cell started with one unit of resource. If a prey organism fed in a given cell (by executing an ‘output’ instruction), the resource in that cell was completely consumed and that cell was replenished with resources over 100 updates.

#### Timing

All Avida trials were run for a set number of updates, as indicated in each of the Phase descriptions below. Updates are the measurement of time in Avida, reflecting the number of genomic instructions executed across the population. Here, each organism executed 30 instructions per update. Organisms had a maximum lifespan of 15,000 instructions executed (500 updates). Realized average number of generations for the populations in each treatment is given in their respective sections below.

#### Reproduction

Organisms, both predator and prey, were required to have consumed 10 resource units and be at least 100 updates old to successfully reproduce. Genome copying and reproduction occurred when the organism executed a single reproduction (‘repro’) instruction. Execution of the reproduction instruction took one update, compared to 1/30th of an update for other instructions (except for predation handling time, as noted below). When an organism reproduced, its offspring was placed in the cell the parent was facing. This new organism’s genome was a copy of the parent’s genome, with the exception any substitutions, insertions, and deletions that occurred during the copy process. Substitutions were independent and only one insertion or deletion was allowed per reproduction (at rates specified in the ‘Phase 1’ and ‘Phase 2’ descriptions, below). In addition to the birth of the new organism upon reproduction, the parent organism was effectively reborn, with all internal states reset. Unlike in legacy Avida experiments, multiple organisms could occupy the same cell [as in 18–20]. Thus, newborn organisms did not replace existing occupants of the cells into which they were born.

#### Predation

In Avida, predation occurs via the evolution of an ‘attack-prey’ instruction, the execution of which allows an organism to attack and kill a non-predator in the cell it is facing. Predation in Avida and the base predator Avida configuration used is described in [17]. Here, as handling time, the attack-prey instruction costs the predator 10 cycles (one-third of an update) if the attack is successful. Predators receive 10% of each prey individual’s consumed resources after a successful kill, which is then applied toward the consumption threshold required for reproduction. An organism is classified as a predator in Avida if it attacks a prey organism, not simply by evolving the ‘attack-prey’ instruction.

In simple (i.e. lacking topographic features like refuges) and confined environments like the one used here, Avida trials containing predators typically require a minimum prey population level below which predator attacks fail. In top-down limited populations like those used here, minimum prey levels prevent population extinctions and serve to standardize prey population sizes and, thus intra-specific competitive pressures. In practice, because prey are constantly being born into populations, prey kills are prevented for very short periods of time and failures simply require predators to make multiple, rapid attacks. Here, attacks on prey were always fatal if there were more than 1,000 prey in the environment (i.e., 1,000 was a minimum prey level). Similarly, in trials without predators, prey population sizes were controlled via a preset maximum population cap (1,000 organisms), as described for each Phase below. When a birth caused that limit to be exceeded, a random prey organism was removed from the population.

### Phase One: Evolutionary History

Our study was divided into three experimental phases (Figure 1). In the first phase (“Phase 1”), we evolved two sets of 30 base populations. Each of these populations was initiated via the placement of 9 identical prey organisms into a new grid environment (Figure 1, Phase 1). These ancestors had simple genomes that allowed them only to blindly move around the environment, repeatedly attempting to feed and reproduce. In one set of populations, the attack-prey instruction was allowed to appear in the organisms’ genomes through random mutations, thus allowing for the potential evolution of predators (Figure 1, Phase 1). Henceforth we refer to these populations as “*predator* EH” treatments. In the second set of populations, the “attack” instruction was prevented from mutating into organisms’ genomes. This second set of “*no predator* EH” base populations thus evolved in the absence of predators in Phase 1. Each population evolved for two million updates (on average, 20,000 generations). During this phase, reproduction carried a 25% probability of a single, random, genomic instruction substitution (maximum one substitution per genome), and 5% probabilities (each) of single insertion and deletion mutations.

**Figure 1.**
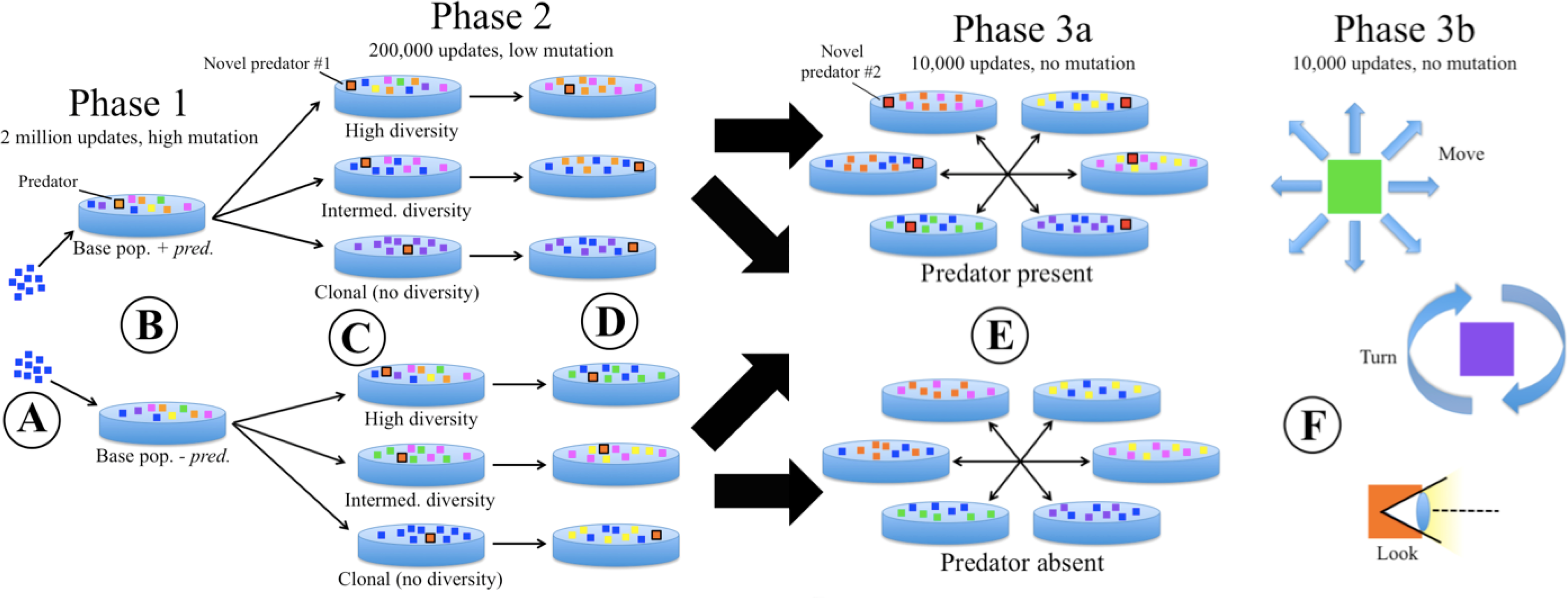
Design of the experiment to test the effects of SGV and EH on evolution of anti-predator traits. A) 9 identical prey are injected to initialize two sets of 30 base populations. B) Over the course of 2 million updates of evolution, populations diversified, including the *de novo* evolution of predators in half of these EH populations (top). C) One fully evolved prey population from each EH treatment (with and without evolving predators) was used to create 30 identical sets of *high*, *intermediate*, and no diversity (*clone*) SGV populations (totaling 2 X 30 X 3 = 180 populations), with clones of the best evolved predator from Phase 1 added to each. D) After a further 2 million updates of evolution at a low mutation rate, replicate populations converged on an intermediate level of diversity, but SGV and EH differences result in trait variation among populations. E) To evaluate the effectiveness of evolved anti-predator traits, each of the 180 fully evolved populations from D were introduced, in turn, into separate competition assays with one randomly selected replicate from each of the other EH X SGV combination populations, both in the presence and in the absence of a new novel predator. F) Separately, prey rates of executing “anti-predator” actions (moves, turns, looks) were recorded in the presence and absence of predators, to quantify expressed behavioral responses to predators for all fully evolved populations.

### Phase Two: Standing Genetic Variation

The second phase (Figure 1, Phase 2) of the experiments allowed us to evaluate how the amount of SGV as well as EH influenced the evolutionary potential of prey in the presence of novel predators. For all Phase 2 trials, prey populations (compositions described below) were evolved for 200,000 updates in the presence of identical copies of a single static predator population (i.e. predators in Phase 2 were prevented from evolving further, dying, or reproducing). Taking in sensory information is a prerequisite for behaviorally responding to and targeting prey, and we used visual input rates as a proxy for predator effectiveness. If organisms evolve to use visual sensors, these sensors provide them with information about their environment, allowing them to “look” at the objects (food, prey or other organisms) in the 45-degree area in front of them. Because predator populations varied across Phase 1 replicates, we selected the Phase 2 predator population from the Phase 1 replicate with highest realized usage of visual sensors (i.e. highest per-predator ‘look-ahead-intercept’ instruction executions) to ensure that an effective predator population was used in Phase 2. Ultimately, the predator population selected and copied for use in all Phase 2 trials had 209 predator organisms.

To create Phase 2 Evolutionary History (EH) treatment source populations, we first selected a single prey population from a random Phase 1 replicate in each EH treatment (*predator* and *no predator* EH), excluding the population from which the Phase 2 predator population was drawn. For each of the two selected populations, we excluded any organisms that were classified as predators or had parents that were predators (as determined by an internal state, see [18,19]), to ensure that Phase 2 prey populations consisted only of prey organisms. An organism must execute an attack instruction to be designated a predator, some prey organisms may have attack instructions in their genomes, provided they have not yet used them; we therefore replaced any attack instructions in the remaining prey populations with a ‘do nothing’ (nop-X) instruction, and prevented any new copies of the attack instruction from mutating into offspring genomes.

To create Phase 2 SGV treatment populations, we created separate “*clone”*, “*intermediate”*, and “*high”* SGV populations by sampling each of the two EH treatment source populations (“seed populations”; one *predator* EH population and one *no predator* EH population: Figure 1, Phase 2). First, for *high* SGV populations, we simply used duplicates of the selected EH treatment source populations. Second, we created each *clone* SGV population by randomly selecting a single genotype from a seed population and making as many duplicates of that genotype as there were organisms in the seed population. A genotype could be selected for multiple Phase 2 clone replicates. Third, we created the *intermediate* SGV populations by randomly sampling genotypes (with replacement) from a seed population, creating up to 55 identical copies of each sampled genotype. We repeated this process until the new *intermediate* SGV population was the same size as the seed population; this sampling scheme yielded *intermediate* SGV prey populations the same size as their *high* SGV seed populations, but with ∼50% lower diversity (Shannon’s diversity index), on average. In all, there were six (2 EH × 3 SGV) Phase 2 treatments, each with 30 replicates.

For each of the six combinations of SGV (*clone, intermediate, high*) and EH (*predator, no predator* EH), we evolved 30 replicate populations for 200,000 updates (500 generations, on average), each in the presence of a copy of the constructed Phase 2 predator population. The mutation rates in Phase 2 were lowered to 0.1% substitution probability, and 0.5% insertion and deletion probability. We found empirically that these mutation rates yielded, on average, 5% divergence (see below) between ancestor and final organisms in Avida after 500 generations (200,000 updates using the Phase 2 settings), matching the expected divergence of a bacterial genome over 500 generations [27]. Other configuration settings were identical to those used in Phase 1.

At the end of Phase 2, we calculated how different the resulting population was from the starting population by using a dynamic programming algorithm [28]. We measured the genetic divergence of each prey organism in each population by aligning the current genome sequence with that of its ancestor from the beginning of Phase 2. Each organism from the seed population was tagged with a unique Lineage ID, which is shared with all progeny of the organism. The Lineage ID of the organisms present in the population at the end of Phase 2 could then be used to identify their ancestor from the beginning of Phase 2. We then calculated the divergence as the percent identity between the two aligned genome sequences. Alignment was required since genome sizes changed during Phase 2, via insertions and deletions.

### Phase 3: Competitive Evaluation of prey populations

For the third and final phase (Figure 1, Phase 3a), we used a set of both “ecological” evaluation simulations to measure the fitness of the final prey populations in the presence and absence of predators, and trait assays to evaluate the evolution of anti-predator traits during Phase 2. For all Phase 3 evaluations, substitution, insertion and deletion mutation rates were set to zero. For all types of Phase 3 trials (described below), in order to create an uneven resource landscape (i.e. in the absence of consumption by prey, all cells would contain resources that could be consumed), populations were introduced into their test environments, run for an initial 1,000 updates, and then reintroduced a second time. Once prey were reintroduced, we ran each trial for an additional 10,000 updates, and recorded population sizes for the two competing populations.

As in Phase 2, Phase 3 trials used copies of a single Phase 1 predator population. The Phase 3 predator was intended to exhibit novel strategies relative to the Phase 2 predator population. We selected the Phase 3 predator population by first eliminating the three Phase 1 replicates used for creating Phase 2 predator and prey populations, and then selecting one predator population that exhibited an average level of visual sensor usage (about half that of the predator population selected for Phase 2). The final selected predator population for Phase 3 contained 253 predators.

Phase 3 involved two sets of trials: the first set allowed for evaluating competitiveness, while the second set allowed for evaluating the effects of SGV and EH on expressed levels of evolved anti-predator behavioral traits. In the first set of Phase 3 trials (Figure 1, Phase 3a), we conducted pairwise competitions in which each population from each of the six Phase 2 treatments was paired, with a randomly selected population from each of the five other treatments (for a total of 30 pairings × 30 replicates = 900 competitions). For each pairing, the two populations were injected once into an environment with the Phase 3 predator population (“*predators present* PT”) and once into an environment without predators (*“predators absent* PT”). For each of these trials, we enforced a prey population level of 2,000 and recorded the relative abundance of each of the two competing populations every 1,000 updates over the course of the 10,000 update trial.

In the second set of Phase 3 trials (Figure 1, Phase 3b), each of the starting (“pre-Phase 2”) and final (“post-Phase 2”) Phase 2 prey populations were combined with a copy of the new Phase 3 predator population and reintroduced into a fresh environment. We then recorded the number of moves and turns executed and the usage of visual sensors (“look” instructions), as proxies for anti-predator responses. Each prey population was then introduced a second time in a separate evaluation in a *predators absent* PT environment. For each of these trials, we kept the number of prey at a constant 1,000 individuals via enforcement of population caps (i.e. a random prey was killed if a new birth would bring the prey population above 1,000) and minimum thresholds (below which predator attacks would fail).

### Statistical analysis

To analyze the effects of SGV, EH, and PT outcomes of ecological competition (Phase 3; Figure 2), we used a linear mixed effects model (Appendix A) with SGV, EH, and PT as fixed effects, and a random intercept and slope across levels of all factors as random effects (Table S3), to account for non-independence of factor levels. For the response variable, a single observation was the mean relative abundance of a single treatment group in a given competition scenario (i.e. one of the two dots in each panel of Figure 2). All first-order interaction terms were included in the model to highlight specific trends. AIC indicated that a second-order interaction did not improve the model (AIC full model: −103.73; AIC reduced model: −112.87; Figure S2).

**Figure 2.**
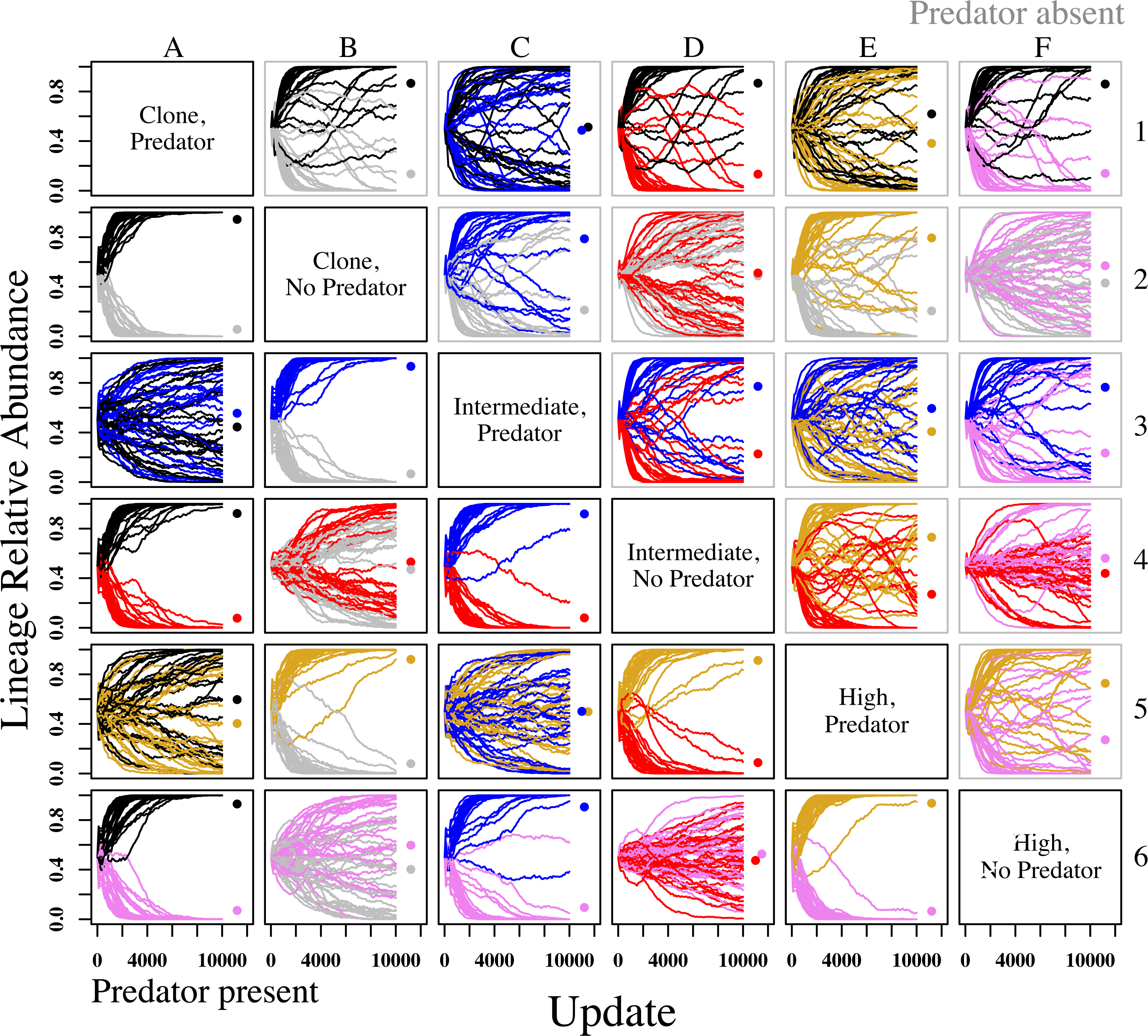
Randomly paired populations in competitions for all SGV and EH treatment combinations in the absence (gray boxes, above diagonal) and presence (black boxes, below diagonal) of a novel predator (Phase 3). EH had a far greater effect on competitive outcomes than SGV in both the presence and absence of predators, with *predator* EH populations competitively dominant in most cases. Y-axis: relative abundance of each lineage in competition. Colors for each lineage indicate SGV and EH treatment history. Black: *clone, predator*; Gray: *clone, no predator*; Blue: *intermediate, predator*; Red: *intermediate, no predator*; Gold: *high, predator*; Pink: *high, no predator*. Dots indicate mean relative abundance for lineages of each treatment after 10,000 updates of competition.

To generate response variables for models describing change in traits due to Phase 2 evolution, we divided instruction counts by total instructions performed (e.g., number of moves / total lifetime instructions) to derive the proportion of instructions represented by each instruction type. Trait assays were conducted pre- and post-Phase 2. Thus, to determine the effect of evolution specifically during Phase 2, we subtracted pre-Phase 2 instruction proportions from post-Phase 2 proportions, yielding the raw change in instruction proportions. All statistical analyses were performed on data scaled by the inverse of the variance to account for heteroscedasticity among treatment groups, though figures in the main text show unscaled data.

We used linear mixed effects models (Equation 1; Appendix B) fit by restricted maximum likelihood to determine the effects of SGV, EH, and Phase 3 predator treatment (PT) on change in total instructions and in instruction proportions due to Phase 2 evolution. SGV, EH, and PT were included as fixed effects. Since both predator treatment (PT) levels (*predators present, predators absent*) were applied to each population, we fitted replicate as a random intercept, and included a random slope across PT levels to account for non-independence of PT levels. Equation 1 describes the change in total prey instructions due to Phase 2 evolution. Models describing change in prey moves, turns, and looks varied in complexity, but were of the same general form (Appendix B). Models describing change in total instructions and changes in proportion moves, proportion turns, and proportion looks were of the same form, but varied in complexity. For each trait, we used a parametrically bootstrapped likelihood ratio test with for model selection, starting with the full model (Equation 1) and dropping interaction terms that did not improve the model fit (Appendix B, Table S4).

#### EQUATION 1

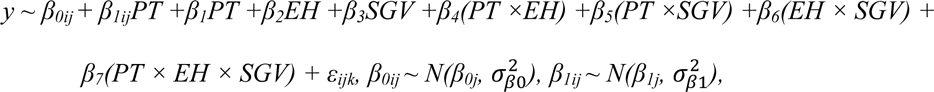

where *β*_*0ij*_ are the random intercepts, and *β*_*1ij*_ are the randomly varying slopes across PT levels. We used a method recently developed by [29] to calculate marginal *R*^2^ (proportion of variation accounted for by fixed effects alone) and conditional *R*^2^ (proportion of variation accounted for by fixed and random effects together). 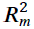 and 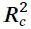 for each model are given in Appendix B, Table S5.

To test the effects of SGV and EH on predator attack rates during trait assays, we used a general linear model with SGV and EH included as fixed effects, and the raw difference in attack rates between pre- and post-Phase 2 evolution as the response variable. A single replicate population was excluded from analyses of change in total instructions, as this replicate had an unusually high pre-Phase 2 instruction count, was highly unrepresentative of prey populations in general, and greatly increased the variance of the *clone* SGV + *no predator* EH treatment group (Figure S7).

### Software & Hardware

We used Avida version 2.14 and the Heads-EX hardware [30] for all experiments. Avida did not originally support immortal, sterile predators as used in Phase 2 and Phase 3, so we added an additional option to Avida that, when set, causes predators to ‘reset’ (akin to being reborn) instead of reproducing or dying of old age. These modifications are available as a patch file against Avida 2.14, along with all analysis scripts, in the GitHub repository (https://github.com/fishjord/avida_predation_scripts).

We conducted all experiments using the Michigan State University’s High Performance Computing Cluster (HPCC). The MSU HPCC contains three general purpose compute clusters containing 2944 cores. Individual Avida runs were submitted as compute jobs to the HPCC general processing queue in parallel where possible. The Avida jobs took between 30 minutes and seven days to run, depending on the number of updates (2,000 to 2,000,000 updates).

We performed all statistical analyses and constructed all figures using the R statistical programming language version 3.0.2. Linear mixed effects models were constructed using the “lmer” function from the “lme4” package (last updated by [31]), and we used the “allEffects” function from the “effects” package (last updated by [32]) to extract marginalized fixed effects. To calculate marginal and conditional R^2^, we used the “r.squaredGLMM” function from the “MuMIn” package (last updated by [33]). We performed bootstrapped likelihood ratio tests using the “PBmodcomp” function from the “pbkrtest” package (last updated by [34]).

## Results

### Ecological competitions

Evolutionary history, i.e. historical exposure to predation in Phase 1, was the most important selective agent shaping competitive abilities of prey both in the presence and in the absence of a novel predator. Prey that evolved with *predator* EH were, in general, stronger competitors than those that evolved with *no predator* EH. In competitions between *predator* EH and *no predator* EH treatments, *predator* EH treatments always had the higher mean final relative abundance (Figure 2; Figure S2 D). Competitive exclusion of a *predator* EH lineage by a *no predator* EH lineage after 10,000 updates in competition was extremely rare (but see e.g. Figure 2, panel D3, in which 1 out of 30 *predator* lineages was excluded). Competitive outcomes were not affected by the presence of a novel predator, regardless of SGV or EH (Figure 2; Figure S2 B, C). While there was no overall effect of SGV on mean final relative abundances, higher SGV seems to have conferred some benefit to *no predator* EH populations (Figure 2; Figure S2 B).

Similarly, there appears to be a subtle interaction between PT and EH, wherein, in the presence of a predator, the difference in relative abundances between predator EH and no predator EH populations during competition assays increases in favor of predator EH populations.

### Evolution of prey traits during Phase 2

Lifetime instruction counts for prey do not reflect higher rates of instruction execution, but rather longer lifespans of prey organisms. Changes in instruction counts varied greatly among treatments (PT × EH × SGV interaction; Figure S3). During Phase 2 evolution, regardless of presence or absence of a novel predator during trait assays, *predator* EH populations generally evolved to increase total instructions executed per lifetime, except in the case of *clone* populations in the absence of predators (Figure 3). Conversely, *no predator* EH populations universally decreased in lifetime instruction counts. In *predator* EH populations, SGV had a markedly positive effect on the magnitude of the decrease in instruction counts, while SGV had little or no effect on instruction counts in *predator* EH populations. Prey generally changed more in total instructions executed with *predators present* than with *predators absent* (Figure 3).

**Figure 3.**
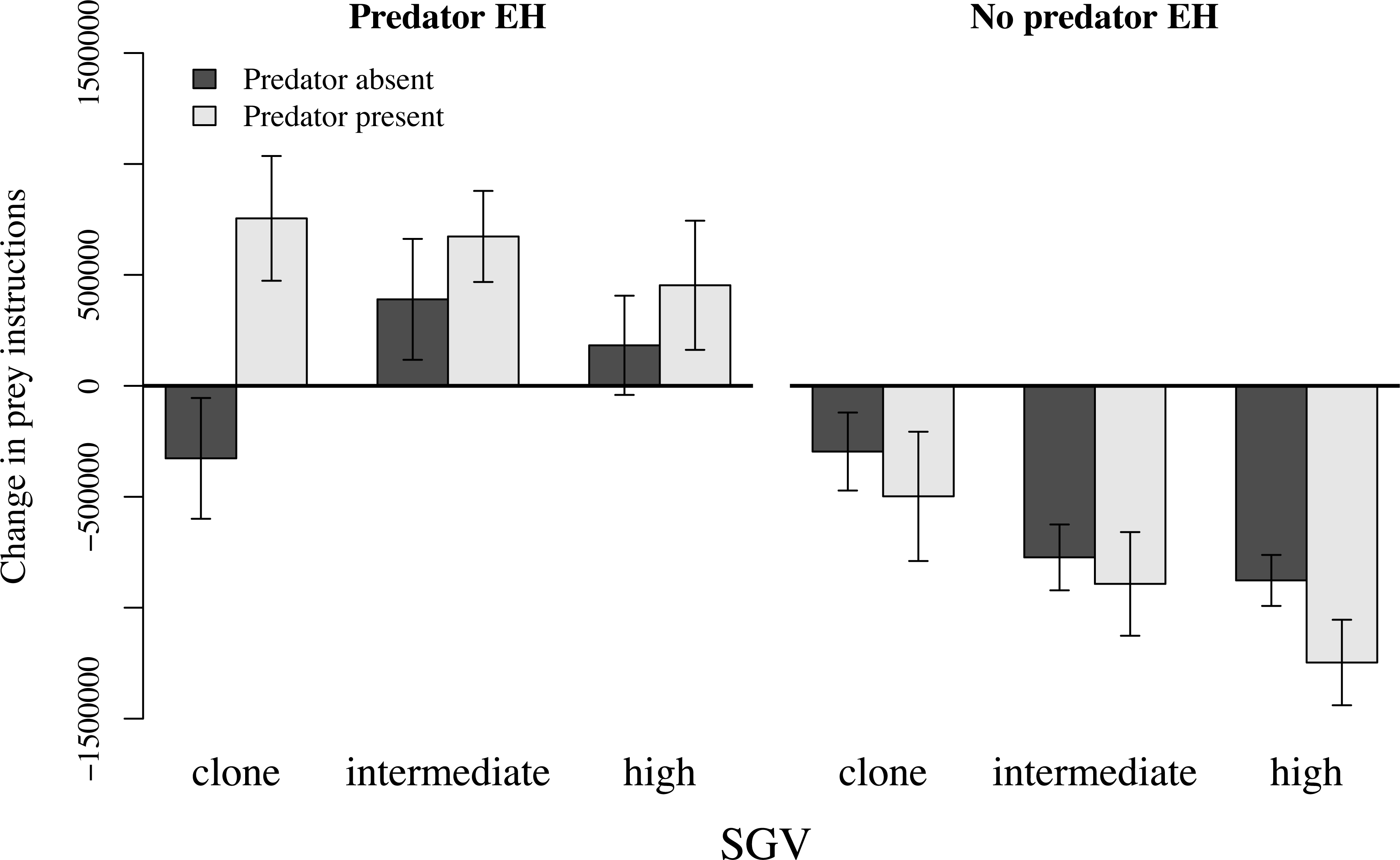
Change in total prey instructions executed (start of Phase 2 minus end of Phase 2), reflecting change in gestation time (or lifespan) resulting from Phase 2 evolution. “*Predator present/absent*” refers to the presence or absence of a novel predator during trait assays before and after Phase 2 evolution. Total instructions either increased or decreased very little in *predator* EH populations, and universally decreased in *no predator* EH populations. Bars are ± 95% CI.

Prey moves as proportions of total prey instructions universally decreased in *predator* EH populations as a result of Phase 2 evolution, and largely increased in *no predator* EH populations (EH effect, Figure 4; Figure S4). Effects of SGV on change in moves were subtle, though slightly less so with *predators absent.* There was no main effect of PT on change in moves (Figure 4; Figure S4).

**Figure 4.**
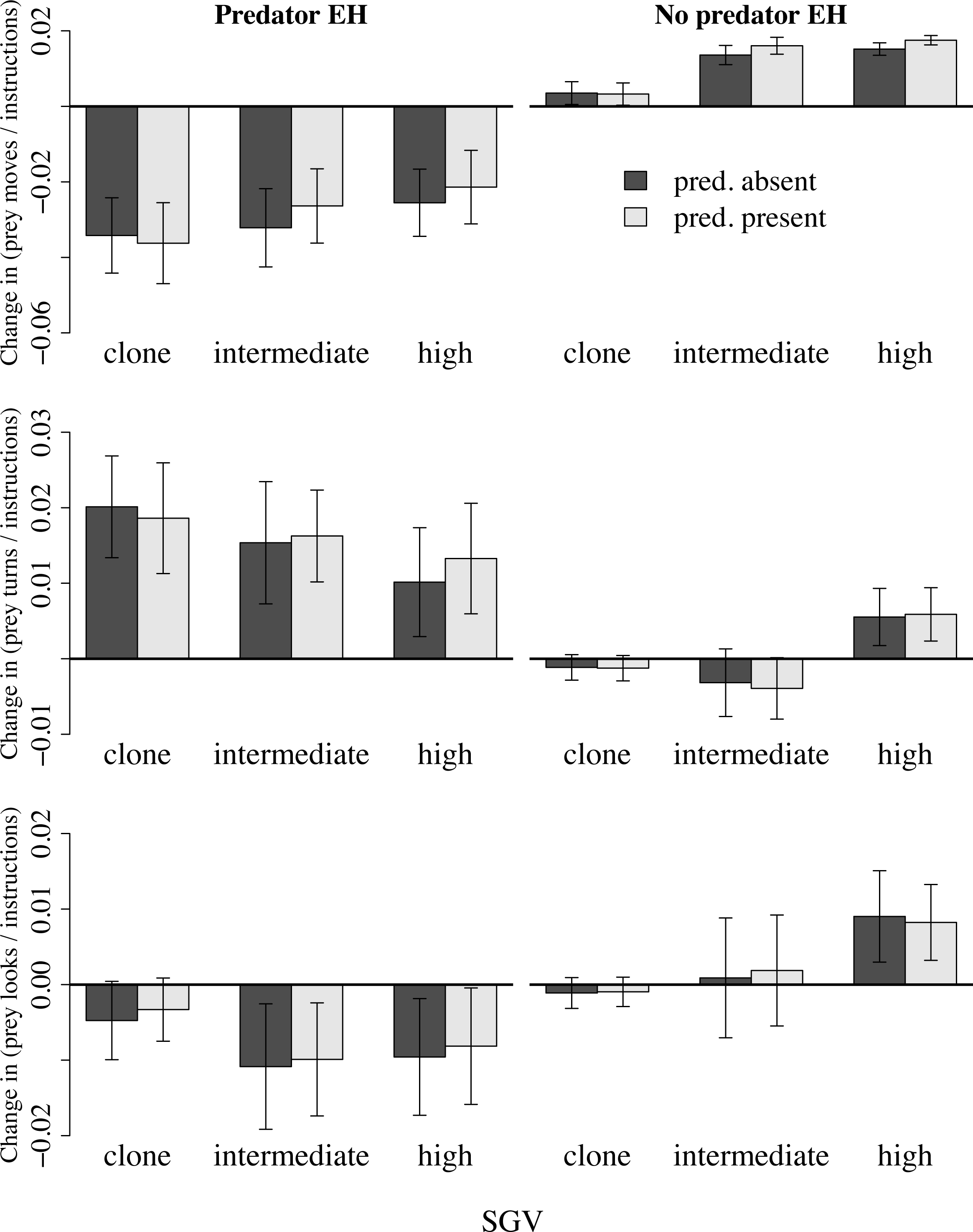
Change in (top) prey moves, (middle) prey turns, and (bottom) prey looks as proportions of total prey instructions, resulting from Phase 2 evolution with a novel predator. “*Predator present/absent*” refers to the presence/absence of the Phase 3 novel predator during the trait assays. Traits in *predator* EH populations generally evolved to a greater extent than did traits in *no predator* EH populations, and traits often evolved in different directions between the two EH treatments. Bars are ± 95% CI.

Prey turns as proportions of total prey instructions increased dramatically in *predator* EH populations, but changed very little in *no predator* EH populations (Figure 4; Figure S5). Effects of SGV were subtle, but were most pronounced in *no predator* EH populations: *clone* and *intermediate* SGV populations decreased slightly on average in proportion of moves, while *high* SGV populations increased (EH × SGV interaction, Figure S5). PT did not affect change in turns (no PT main effect, Figure 4; Figure S5), nor was there a main effect of SGV (Figure S5).

Proportions of instructions that were looks generally decreased in *predator* EH populations, but changed little in *no predator* EH populations; EH effects varied among levels of SGV, with the smallest EH effects in *clone* SGV populations (EH × SGV interaction, Figure 4; Figure S6). Main effects of SGV and PT were not statistically significant (Figure S6).

### Effects of Phase 2 prey evolution on predator attack rates

Phase 2 evolution led to a reduction in predator attacks in nearly all cases, and evolutionary history and standing genetic variation both affected predator attacks on prey (Figure 5; Table S2). *High* and *intermediate* SGV populations experienced qualitatively similar reductions in predator attacks as a result of Phase 2 evolution, and prey evolved with *predator* EH experienced similar reductions in predator attacks across SGV treatments (Figure 5). Only *clone* SGV + *no predator* EH populations were attacked more after Phase 2 evolution.

**Figure 5.**
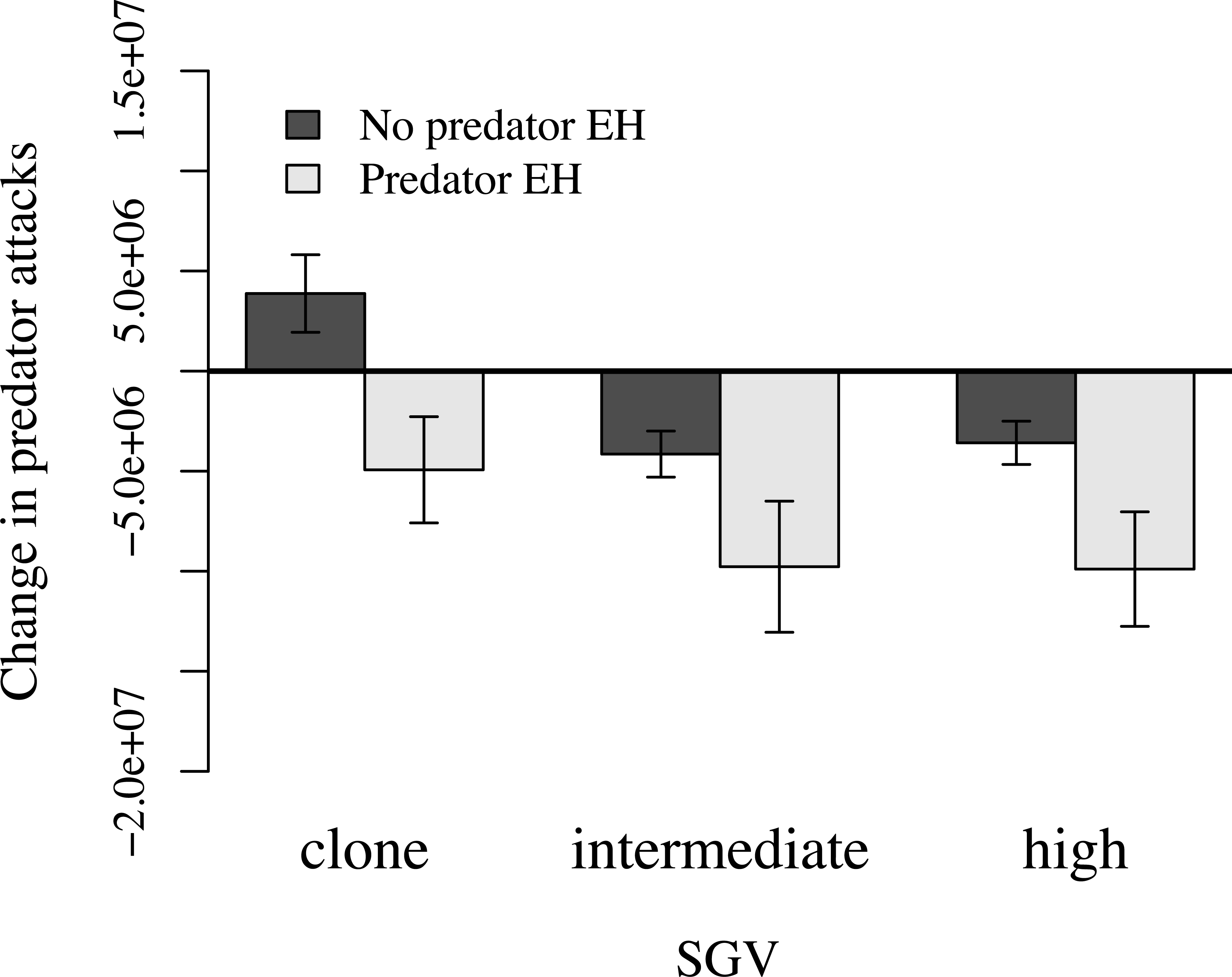
Change in attacks in behavioral assays pairing the Phase 3 predator with pre- and post- Phase 2 prey populations. Predator attack frequencies on *predator* EH populations universally decreased, and generally changed more than did attack frequencies on *no predator* EH populations. Bars are ± 95% CI.

## Discussion

Though the effects of evolutionary history (EH) and standing genetic variation (SGV) on the evolutionary potential of populations have been the subject of considerable conjecture [5,9], they have rarely been tested. Here, we were able to separately control for and vary the background levels of SGV and EH, and subsequently test the roles of each in shaping the evolvability of populations in a predator-prey context.

### Role of evolutionary history

Evolutionary history (the combined selective pressures under which populations evolve) likely determines the nature of mutations that fix in populations [9]. We found that having an evolutionary history with a predator improves a population’s ability to evolve defenses in response to novel pressures from novel invading predator populations: in general, *predator* EH populations experienced greater changes in trait expression compared to *no predator* EH populations (Figure 4) when evolving defenses against invading (Phase 2) predators. This occurs regardless of the level of SGV, suggesting that the importance of EH eclipses that of evolvability originating from SGV. Furthermore, in pairwise competitions (Figure 2), populations with *predator* EH demonstrated competitive superiority over *no predator* EH populations, similar to findings of [18]. If a novel predator was present during the competition, the advantage was even more pronounced (also see Figure S2 C), with the vast majority of *predator* EH populations competitively excluding their *no predator* EH competitors (Figure 2). Additionally, when introduced into separate ecological trials with the novel predator, populations that evolved with *predator* EH suffered fewer attacks than populations evolved with *no predator* EH (Figure 5).

Because both types of EH populations were further evolved in the presence of predators (in Phase 2) before they were competed against each other, the ecological advantage enjoyed by *predator* EH populations in competition with *no predator* EH populations was due, in part, to their historic evolution of anti-predator traits (i.e. in Phase 1), even though that initial evolution was in response to an entirely different predator population.

Both the mutation rates used for the evolution of phase 1 predator EH populations and the duration of the evolutionary trials will have influenced the extent of evolutionary history in each trial (i.e. the degree to which prey populations were adapted to predators). Because we did not vary Phase 1 mutation rates or duration, we cannot comment on how experimentally altering the strength of evolutionary history could impact the observed trends.

### Role of standing genetic variation

Many studies have pointed to SGV as a key component of a population’s evolutionary potential [6]. When there is greater genetic variation in a population, there is more raw material upon which natural selection can act. Thus, we expected SGV alone to play a significant role in the rapid evolution of anti-predator traits. However, we found only small effects of historic SGV in determining the final outcomes of ecological competitions (Figure 2). Furthermore, the effects of SGV on the evolution of specific anti-predator traits (moves, turns, looks) were detectable only within EH treatments (Figure 4).

### Standing genetic variation (SGV) vs Evolutionary history (EH)

Across the *predator* EH populations, we found no detectable effects of SGV on anti-predator trait values (Figure 4) or competitive outcomes (Figure 2). We suggest that a lack of an SGV effect on *predator* EH populations may be due, in part, to the historical predator weeding out unfit prey genotypes. E.g., predators reduced prey diversity (Figure S1; Table S1) because they operate as a very strong agent of selection against less fit phenotypes. However, we would expect SGV to have an effect within *no predator* EH populations (as seen in Figure 4). Given the lack of an evolutionary history with predators, in *no predator* EH populations, the odds of one sampled genotype (*clone* SGV) being able to quickly acquire a beneficial mutation are low. On the other hand, if the full suite of all discovered genotypes (*high* SGV) remains in the population, there is a greater potential for providing precursory material for rapidly generating anti-predator traits. However, if EH leads to a large number of genotypes having the prerequisites for rapid adaptive response, the odds of sampling a beneficial genotype are high, and differences in SGV will not significantly affect evolvability.

Indeed, as we have shown, in order to adaptively address novel predation pressures, it appears to be easier to modify historically realized and relevant adaptations than to repurpose unrelated genes, or evolve effective traits *de novo*. Thus, the greater evolvability of the *predator* EH populations would have arisen out of their ability to adapt anti-predator strategies across predator contexts. More broadly, evolvability may commonly be determined by a population’s ability to use strategies across environmental contexts, i.e. their adaptive complexity [35].

## Acknowledgements

This work was supported by the BEACON Center for the Study of Evolution in Action (NSF Cooperative Agreement DBI-0939454) and the Michigan State University Institute for Cyber Enabled Research. We thank Chris Adami for comments and feedback on an earlier draft of this manuscript, and Max McKinnon for contributions to early stages of this research.

---

**Figure S1.**
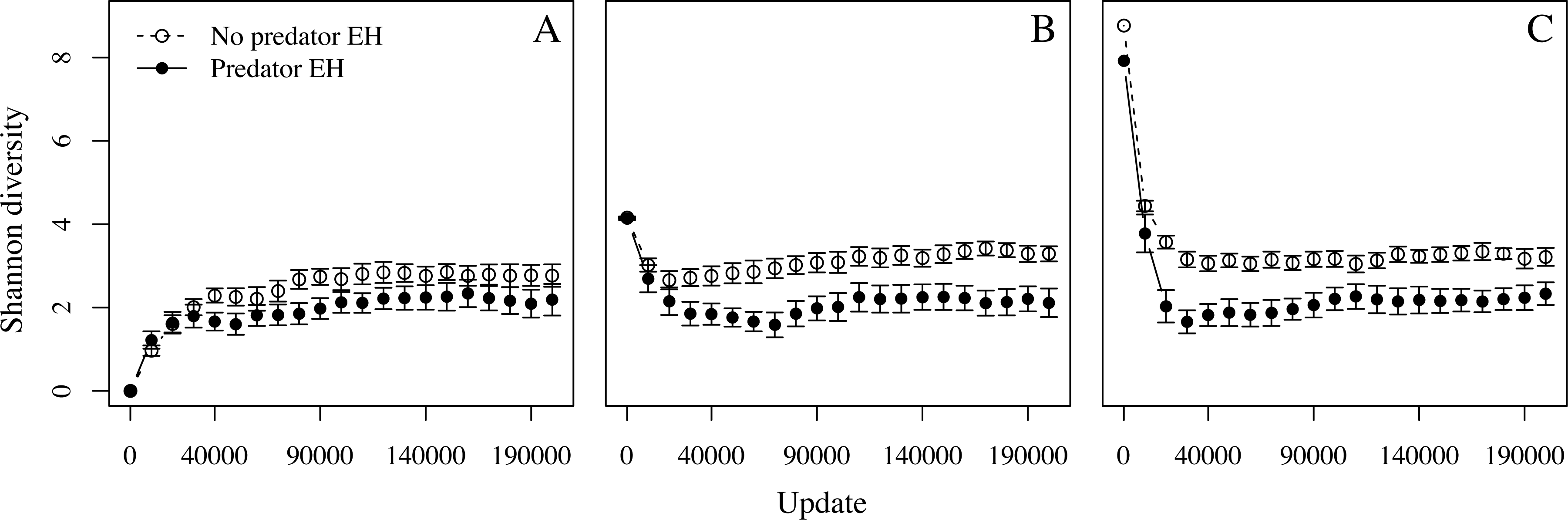
Shannon’s diversity across 200,000 updates of Phase 2 evolution in (A) *high*, (B) *intermediate*, and (C) *clone* SGV treatments. Filled circles represent *no predator* EH treatments, open circles represent *predator* EH treatments. Bars are ± 95% CI.

**Figure S2.**
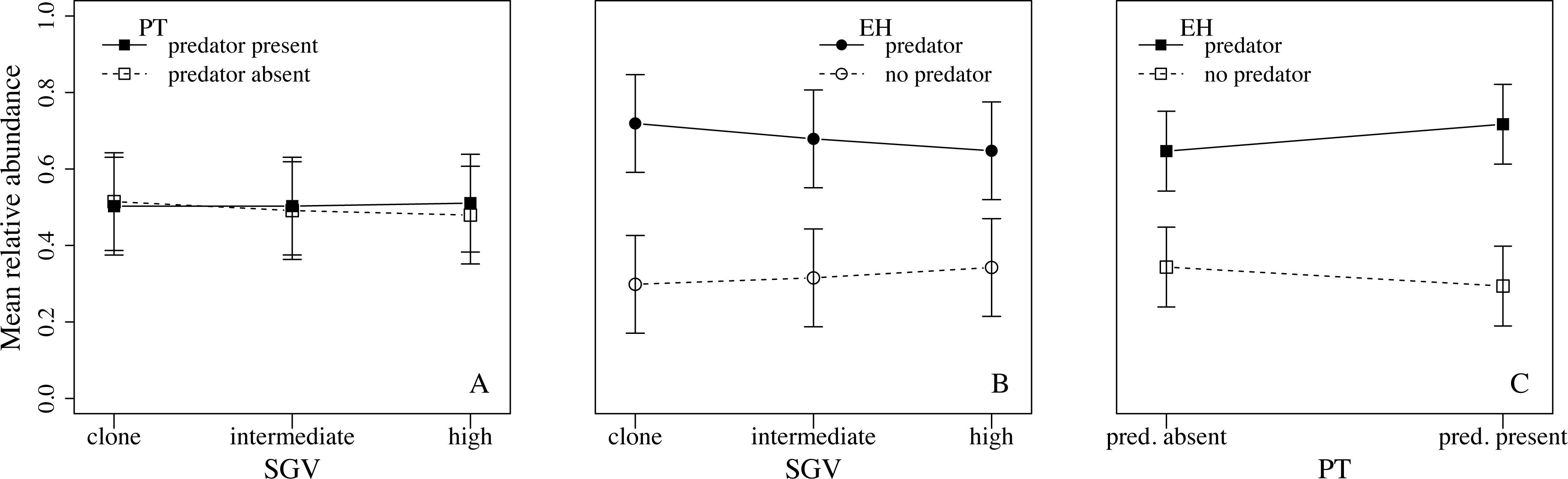
Marginalized fixed effects of a generalized linear mixed effects model describing effects of SGV, EH, and PT on mean relative abundance of a single treatment group from each competition scenario depicted in Figure 2. **A**. SGV × PT interaction plot; **B**. SGV × EH interaction plot; **C**. EH × PT interaction plot. Bars are ± 95% CI. EH was by far the most important factor explaining competitive outcomes; *predator* EH groups were universally superior to *no predator* EH groups in ecological competition. Formal model, random effects output, and marginal and conditional R^2^ can be found in Appendix A.

**Figure S3.**
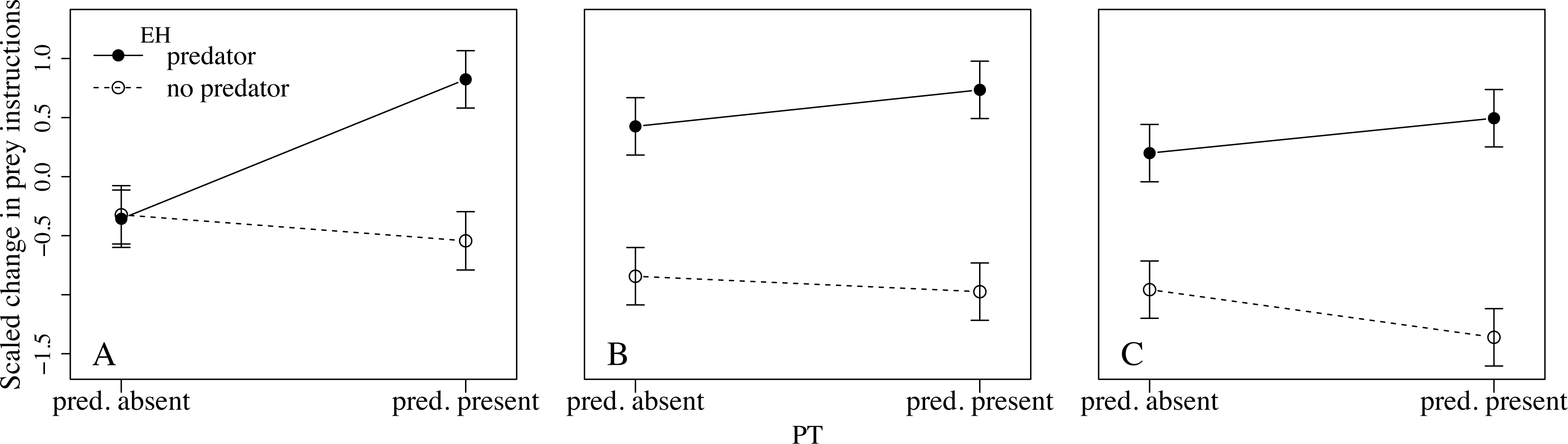
Marginalized fixed effects of a generalized linear mixed effects model of scaled change in total prey instructions across Phase 2 evolution as a function of SGV (**A**. clone; **B**. intermediate; **C**. high), EH, and PT. Bars are ± 95% CI. EH had the greatest effect on change in prey instructions, though PT also affected change in instructions, especially in *clone* SGV populations. Formal model, random effects output, and marginal and conditional R^2^ can be found in Appendix B.

**Figure S4.**
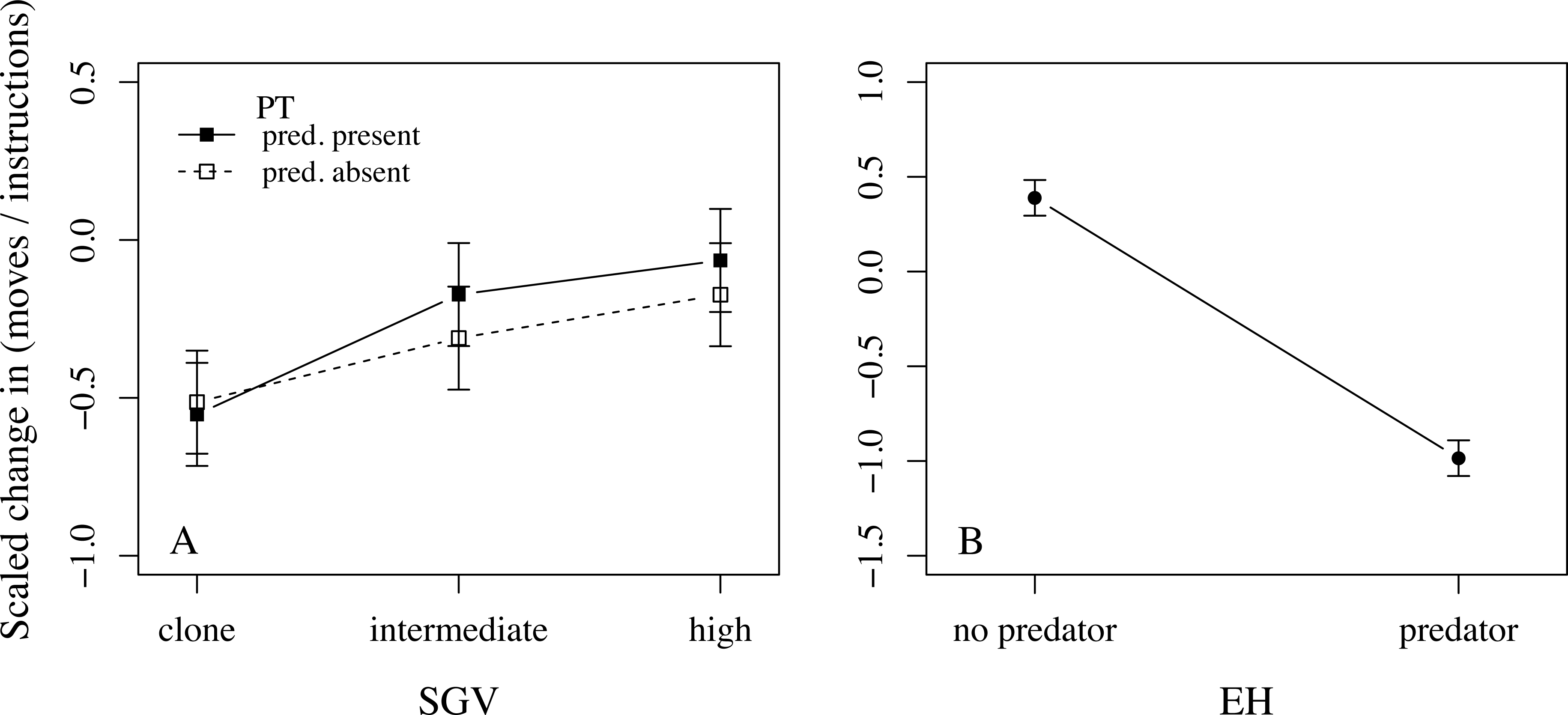
Marginalized fixed effects of a generalized linear mixed effects model of scaled change across Phase 2 evolution in the proportion of prey instructions that were moves as a function of **A**. SGV and PT and **B**. EH. Bars are ± 95% CI. Both EH and SGV affected change in proportion moves, with SGV increasing more dramatically across SGV treatments with *predator present*. Formal model, random effects output, and marginal and conditional R^2^ can be found in Appendix B.

**Figure S5.**
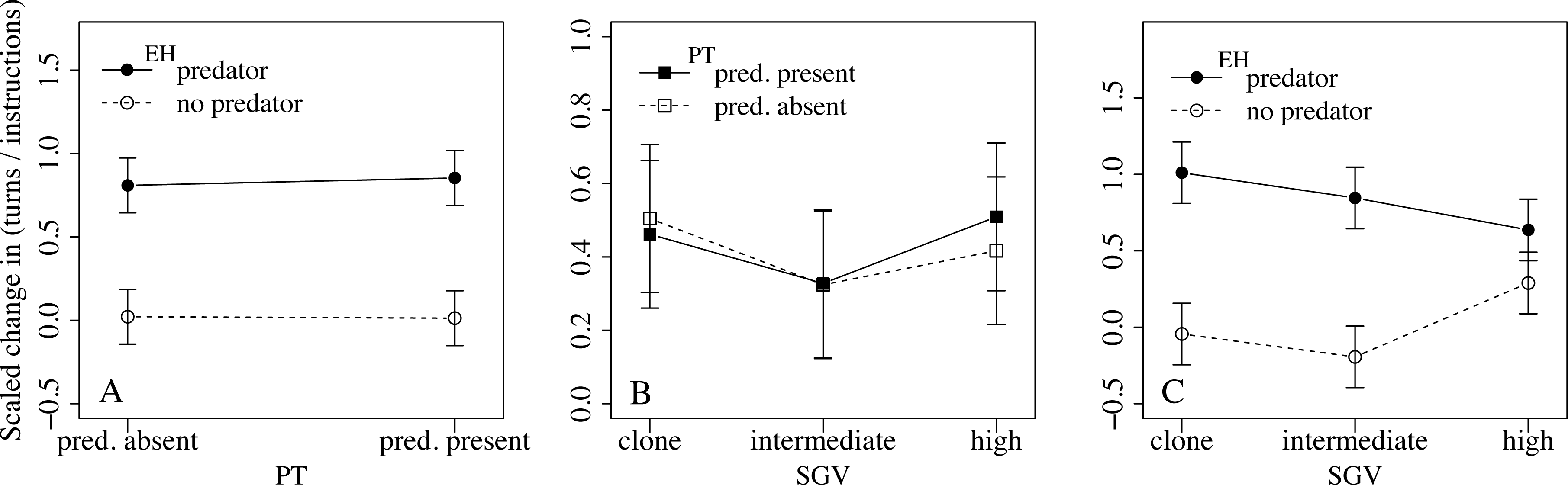
Marginalized fixed effects of a generalized linear mixed effects model of scaled change across Phase 2 evolution in the proportion of prey instructions that were turns as a function of **A**. PT and **B**. SGV × EH. Bars are ± 95% CI. *Predator* EH populations increased turns as a results of Phase 2 evolution; change in turns was marginal for *no predator* EH populations, and SGV affected *predator* and *no predator* populations differently. PT had no detectable effect. Formal model, random effects output, and marginal and conditional R^2^ can be found in Appendix B.

**Figure S6.**
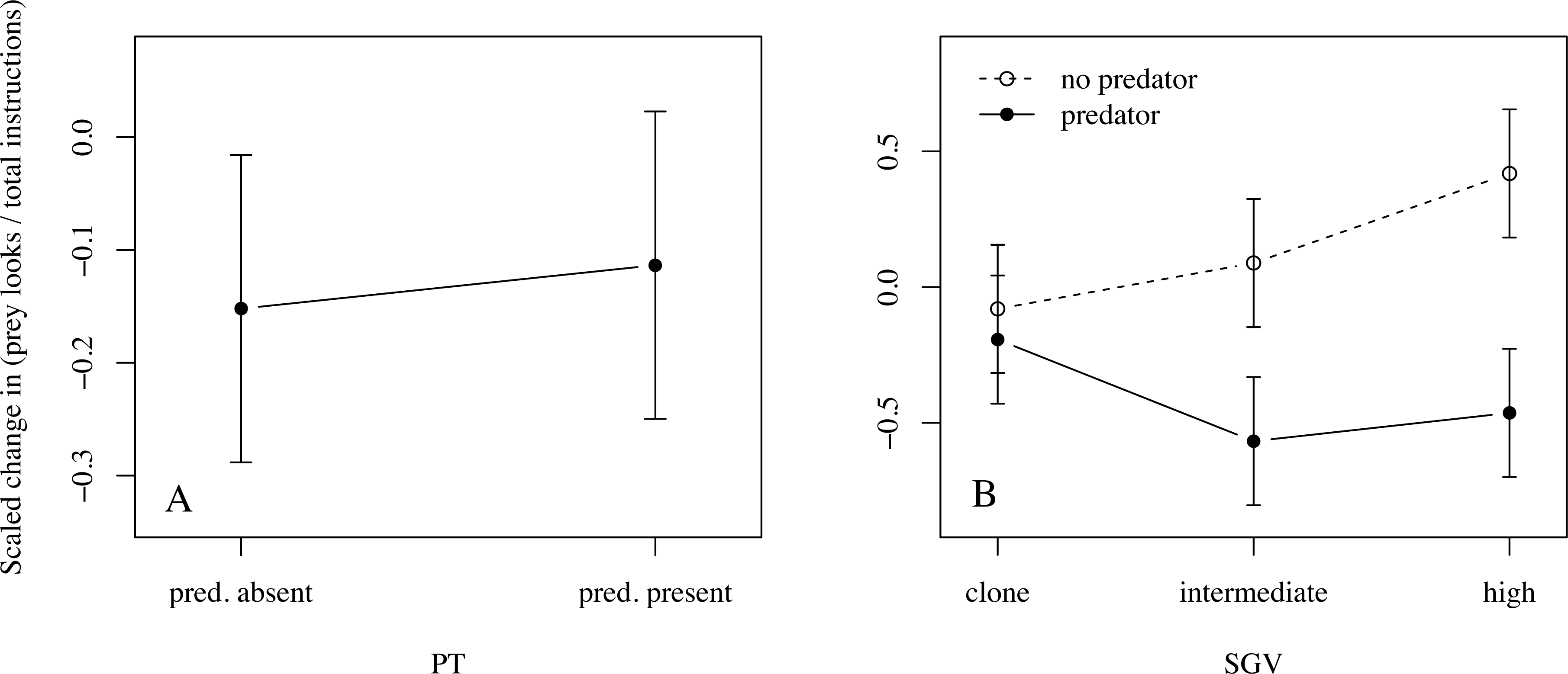
Marginalized fixed effects of a generalized linear mixed effects model of scaled change across Phase 2 evolution in the proportion of prey instructions that were looks as a function of **A**. PT and **B**. SGV × EH. Bars are ± 95% CI. *Clone* and *high* SGV populations with *predator* EH increased looks as a result of Phase 2 evolution and change in turns was marginal to negative for *no predator* EH populations. PT had no detectable effect. Formal model, random effects output, and marginal and conditional R^2^ can be found in Appendix B.

**Figure S7.**
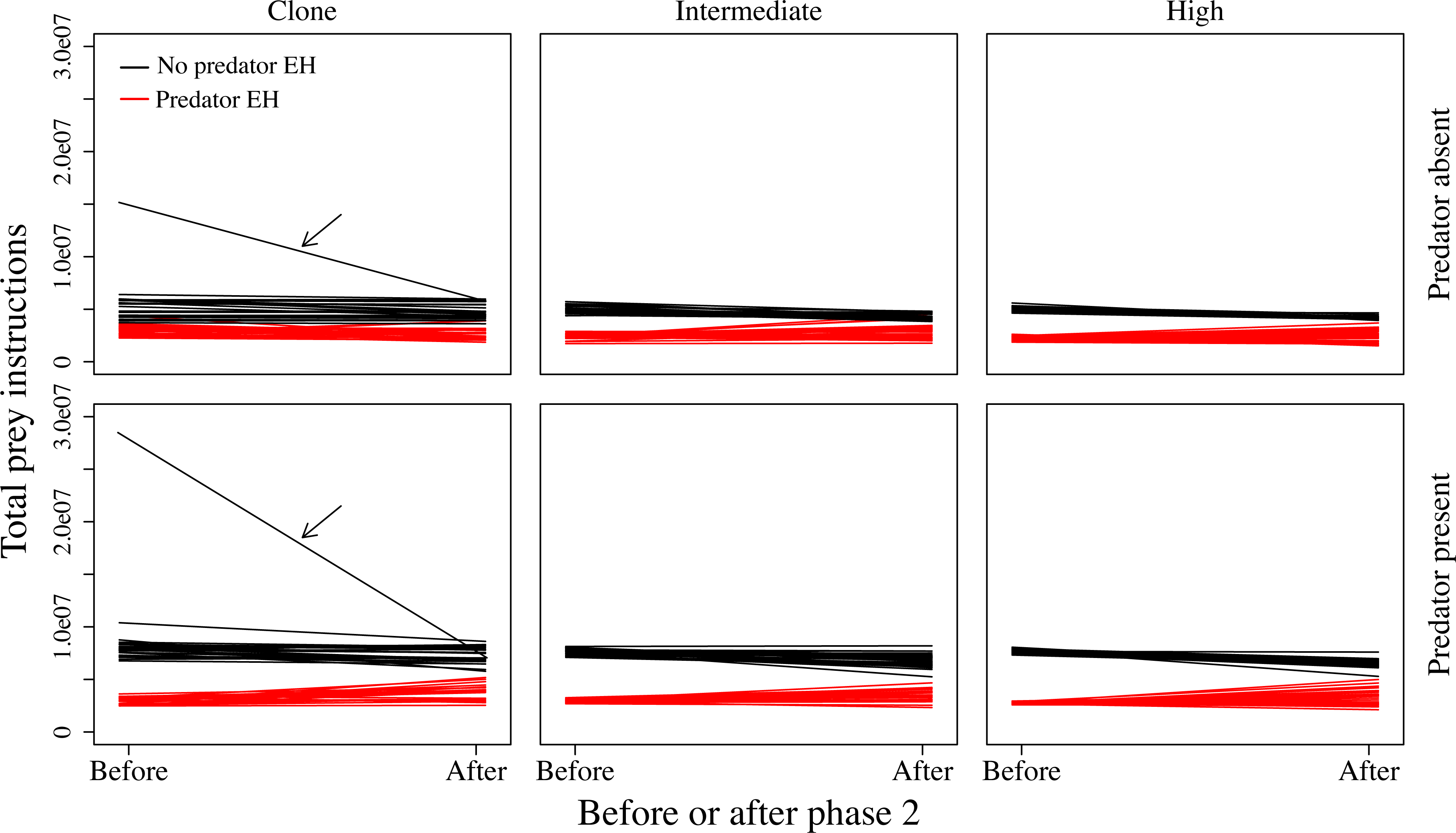
Total instructions executed by each replicate prey population before and after Phase 2 evolution. The slope of each line represents the evolutionary change in instructions executed per lifetime (i.e. gestation time). The same replicate populations were assayed in the absence (top) and in the presence (bottom) of the Phase 3 predator. Arrows indicate the replicate population that was excluded from all statistical analyses, as it is not representative of its treatment group (*clone* SGV, *no predator* EH).

## Supporting Information (S1)

### Genetic diversification during Phase 2 evolution

Over the 200,000 updates of Phase 2 evolution, *clone* SGV populations increased in diversity (Shannon’s Diversity Index), and *intermediate* and *high* SGV populations decreased in diversity, with all trends stabilizing near the 40,000^th^ update (Figure S1). Both standing genetic variation (SGV) and evolutionary history (EH) affected diversity values at the end of Phase 2 evolution. Diversity was similar across SGV treatments for *predator* EH populations, but was higher in *intermediate* and *high* than in *low* SGV populations for *no predator* EH populations. Shannon’s Diversity was higher over all for *no predator* than for *predator* EH populations (Figure S1; Table S1).

### Supplemental tables (S2)

**Table S1.**
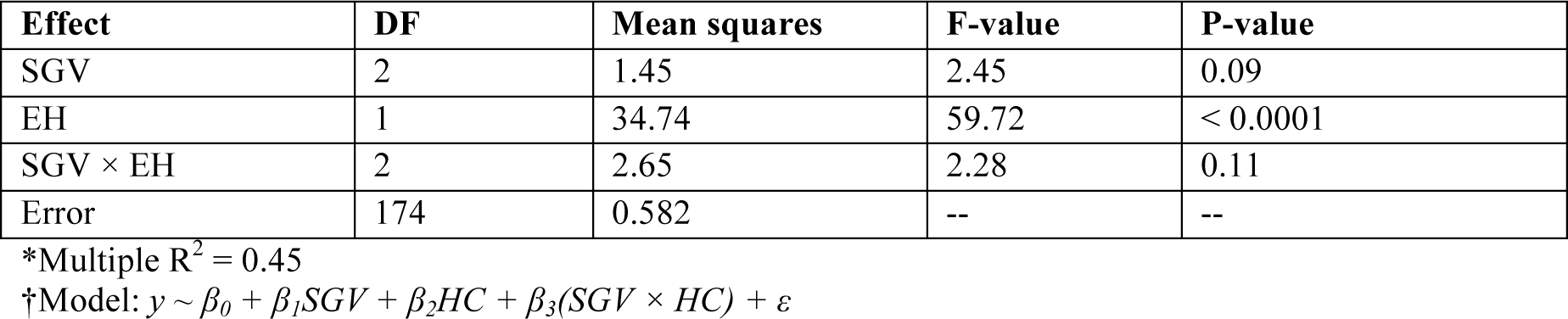
Analysis of Variance (calculated using the anova() function in the R 3.0.2 base package) on a linear model*† testing the effects of SGV and EH and their interaction on Shannon’s diversity among populations at the 200,000^th^ update of Phase 2 evolution.

**Table S2.** Analysis of Variance (calculated using the anova() function in the R 3.0.2 base package) on a linear model*† testing the effects of SGV and EH and their interaction on change in predator attacks as a result of Phase 2 evolution.

| Fixed effect | DF | Mean squares | F-value | P-value |
| --- | --- | --- | --- | --- |
| SGV | 2 | $1.60 \times 10^{15}$ | 19.89 | < 0.0001 |
| EH | 1 | $4.30 \times 10^{15}$ | 53.51 | < 0.0001 |
| SGV × EH | 2 | $8.41 \times 10^{13}$ | 1.05 | 0.35 |
| Error | 354 | $8.04 \times 10^{13}$ | -- | -- |

\*Multiple $R^2 = 0.20$
†Model: $y \sim \beta_0 + \beta_1SGV + \beta_2EH + \beta_3(SGV \times EH) + \varepsilon$

### Appendix A (S3)

Linear mixed effects model describing effects of SGV, EH, and PT on mean final relative abundances of treatment groups in pairwise ecological competitions (Figure 2; Figure S2).

Model A.1:

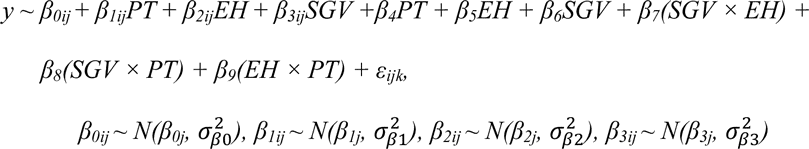

*β*_*0ij*_ are the randomly varying intercepts, *β*_*1ij*_ are the randomly varying slopes across PT levels, *β*_*2ij*_ are the randomly varying slopes across EH levels, and *β*_*3ij*_ are the randomly varying slopes across SGV levels.

**Table S3.** Random intercepts and slopes (linear contrasts) fit in Model A.1 with parametric bootstrap 95% confidence intervals.* Different replicate ID’s were assigned by treatment combination for each factor. 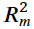 = 0.51; 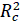 = 1.0.

| Factor | Coefficient | Std. dev. | Lower 95% | Upper 95% |
| --- | --- | --- | --- | --- |
| SGV |  |  |  |  |
|  | slope (intermediate / clone) | 0.030 | 0.030 | 0.030 |
|  | slope (high / clone) | 0.030 | 0.030 | 0.030 |
|  | intercept | 0.18 | 0.17 | 0.18 |
| *EH* |  |  |  |  |
|  | slope | 0.038 | 0.038 | 0.039 |
|  | intercept | 0.031 | 0.030 | 0.031 |
| *PT* |  |  |  |  |
|  | slope | 0.015 | 0.015 | 0.015 |
|  | intercept | 0.0088 | 0.0087 | 0.0089 |
| Residuals | -- | 0.0018 | 0.0017 | 0.0018 |

\*Some confidence interval values are the same as standard deviations, due to rounding.

### Appendix B (S4)

Linear mixed effects models describing change in prey instructions across Phase 2, compared using LRT with a parametric bootstrap. Models are nested, with Model 1 being the full model (all first and second-order interactions included), and Model 6 being the simplest (no interactions included). Models 2–5 include some (Models 3–5) or all (Model 2)"first-order interactions. Interaction terms included were those determined to be statistically significant (parametric bootstrap confidence intervals around coefficients did not overlap zero). LRT outputs, AIC for each model, and model selection processes are given in Table S3.

Model B.1 (full model):

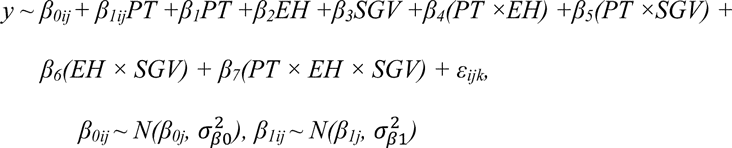

Model B.2:

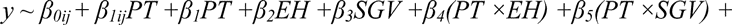

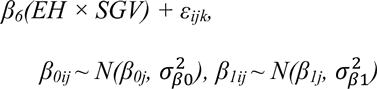

Model B.3:

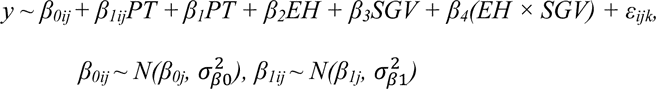

Model B.4:

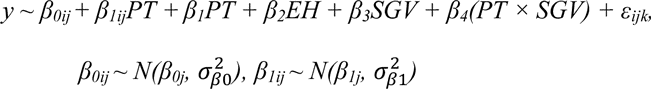

Model B.5 (fully reduced model):

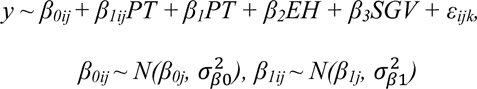

*β*_*0ij*_ are the randomly varying intercepts, and *β*_*1ij*_ are the randomly varying slopes across PT levels.

**Table S4.** AIC, LRT output, and model selection processes for linear mixed-effects models describing change in **A**. total prey instructions; **B**. proportion moves; **C**. proportion turns and **D**. proportion looks.

| Compared | AIC |  | X<sup>2</sup> | df | LRT P-val | PBtest P-val | Mod. preferred |
| --- | --- | --- | --- | --- | --- | --- | --- |
| A. Total instructions |  |  |  |  |  |  |  |
| B.1 & B.2 | 678.09 | 692.23 | 18.14 | 2 | 0.0001 | 0.001 | 1, 1 chosen |
| B. Proportion moves |  |  |  |  |  |  |  |
| B.1 & B. 2 | 370.83 | 370.82 | 3.98 | 2 | 0.14 | 0.15 | none, 2 chosen |
| B.2 & B.4 | 370.82 | 367.76 | 1.99 | 3 | 0.57 | 0.55 | none, 4 chosen* |
| C. Proportion turns |  |  |  |  |  |  |  |
| B.1 & B.2 | 587.37 | 586.37 | 3.98 | 2 | 0.14 | 0.16 | none, 2 chosen |
| B.2 & B.4 | 586.37 | 589.67 | 9.30 | 3 | 0.03 | 0.03 | 2, 2 chosen |
| *D. Proportion looks* |  |  |  |  |  |  |  |
| B.1 & B.2 | 598.84 | 595.98 | 1.14 | 2 | 0.57 | 0.59 | none, 2 chosen |
| B.2 & B.3 | 595.98 | 592.18 | 2.20 | 3 | 0.53 | 0.53 | none, 3 chosen |
| B.3 & B.5 | 592.18 | 594.18 | 6.00 | 2 | 0.05 | 0.05 | 3, 3 chosen |

\*All remaining first-order interactions significant, so no further model comparison.

**Table S5.** Random intercepts and slopes (linear contrasts) between levels of PT with parametric bootstrap 95% confidence intervals for models describing change in **A**. total prey instructions; **B**. proportion moves; **C**. proportion turns; and **D**. proportion looks. Also included are marginal and conditional R^2^ and residual standard deviations with parametric bootstrap 95% CI.

| Trait | Coefficient | Std. dev. | Lower 95% | Upper 95% |
| --- | --- | --- | --- | --- |
| *A. Total instructions: $R_m^2 = 0.57$; $R_c^2 = 0.82$* |  |  |  |  |
|  | slope | 0.46 | 0.38 | 0.52 |
|  | intercept | 0.29 | 0.21 | 0.35 |
|  | residuals | 0.39 | 0.34 | 0.41 |
| *B. Proportion moves: $R_m^2 = 0.56$; $R_c^2 = 0.97$* |  |  |  |  |
|  | slope | 0.61 | 0.54 | 0.67 |
|  | intercept | 0.02 | 0.004 | 0.06 |
|  | residuals | 0.17 | 0.15 | 0.19 |
| *C. Proportion turns: $R_m^2 = 0.23$; $R_c^2 = 1.0$* |  |  |  |  |
|  | slope | 0.81 | 0.72 | 0.88 |
|  | intercept | 0.37 | 0.32 | 0.40 |
|  | residuals | 0.00044 | 0.00040 | 0.00048 |
| *D. Proportion looks: $R_m^2 = 0.10$; $R_c^2 = 1.0$* |  |  |  |  |
|  | slope | 0.96 | 0.83 | 1.03 |
|  | intercept | 0.27 | 0.23 | 0.31 |
|  | residuals | 0.13 | 0.12 | 0.14 |

